# In airway epithelium, basal stem cells and their stress fibers remodel during the unjamming transition

**DOI:** 10.1101/2022.08.18.504453

**Authors:** Thien-Khoi N. Phung, Jennifer A. Mitchel, Michael J. O’Sullivan, Jin-Ah Park

**Author notes:** **Corresponding author:** Jin-Ah Park, Ph.D., Program in Molecular and Integrative Physiological Sciences, Department of Environmental Health, Harvard T.H. Chan School of Public Health, 665 Huntington Ave, Rm1-315, Boston, Massachusetts, USA.

## Abstract

Under homeostatic conditions, epithelial cells remain non-migratory. However, during embryonic developmental and pathological processes, they become migratory. The mechanism underlying the transition between non-migratory and migratory epithelial cells is a fundamental question of cellular biology. In well-differentiated primary human bronchial epithelial cell layers, non-migratory epithelial cells become migratory through an unjamming transition (UJT). We have previously identified the hallmarks of UJT: apical cell elongation and collective cellular migration. These indicate that UJT is driven by intercellular force modulation, but the nature of these forces in pseudostratified epithelia is unknown. Here, we identify structural characteristics of basal stem cells that are indicative of force generation. During the UJT, basal stem cells elongate and enlarge, and their stress fibers lengthen and align. These morphological changes in basal stem cells correspond to the previously defined hallmarks of the UJT. Moreover, basal cell elongation and stress fiber lengthening precedes apical cell elongation. Together, these structural changes in basal stem cells suggest that in pseudostratified airway epithelium, basal stem cells may be the origin of the traction forces through stress fiber modeling during the UJT.

**Summary Statement:** Our image analysis of pseudostratified airway epithelium reveals basal stem cells as the likely source of traction forces driving collective cellular migration during an unjamming transition.

## Introduction

Epithelial cells lining the large airways are complex in their differentiated cellular composition which forms a pseudostratified columnar epithelium (Montoro et al., 2020). The pseudostratified human airway epithelium can be recapitulated *in vitro* by culturing airway basal stem cells from the large airway in air-liquid interface (ALI) culture (Gray et al., 1996; O’Sullivan et al., 2021). During ALI culture, epithelial layer maturation is marked by a decrease in cellular migration speed across the epithelial layer (Park et al., 2015). As the basal stem cells progressively differentiate through ALI days, the confluent layer transitions from a migratory unjammed phase towards a sedentary jammed phase; this process is termed the jamming transition. When this jammed layer is exposed to pathologic stimuli, such as mechanical compression (mimicking bronchospasm) or ionizing radiation, epithelial cells become migratory again. Our previous work demonstrated that this process of gaining motility in a pseudostratified epithelium takes place through an unjamming transition (UJT) (Mitchel et al., 2020; O’Sullivan et al., 2020; Park et al., 2015). Unlike the partial epithelial-to-mesenchymal transition (pEMT), a well-studied mechanism for achieving collective cellular migration, during the UJT the epithelium maintains its pseudostratified structure and barrier integrity (Mitchel et al., 2020). Epithelial migration is critical for physiological and pathologic conditions, including embryonic development, cancer metastasis, and tissue repair. In particular, the UJT has been implicated in airway branching morphogenesis as well as in chronic lung diseases (Park et al., 2015; Spurlin et al., 2019; Stancil et al., 2021; Stancil et al., 2022). However, the underlying physical mechanism for collective migration during UJT in pseudostratified epithelia remains unknown.

For collective cellular migration in a simple columnar epithelium, cells generate traction forces on their underlying substrate through remodeling of their actin stress fibers. These forces are transmitted between neighboring cells and across the epithelial layer predominantly in the plane of migration (Kim et al., 2013; Tambe et al., 2011; Trepat et al., 2009). These intercellular forces drive collective cellular migration and are measurable using traction force microscopy and monolayer stress microscopy (Butler et al., 2002; DeCamp et al., 2020; Saraswathibhatla and Notbohm, 2020; Saraswathibhatla et al., 2020; Tambe et al., 2013; Trepat et al., 2009). For example, Saraswathibhatla and colleagues demonstrated that traction forces measured by traction force microscopy are increased during the UJT (Saraswathibhatla and Notbohm, 2020; Saraswathibhatla et al., 2021). Moreover, they showed increased traction force is associated with alignment of stress fibers. However, the traction and intercellular forces measured in a simple columnar epithelium cannot be intuitively extrapolated for a pseudostratified epithelium. The pseudostratified epithelium, such as that found in the airway, is composed of basal stem cells positioned underneath well-differentiated luminal cells. These well-differentiated cells project mainly to the apical surface, but are believed to have a relatively small footprint connecting to the underlying basement membrane (Evans et al., 2001). Therefore, during collective migration interactions between the apical and basal cells in pseudostratified epithelia remain challenging to decipher. In addition, studies from simple columnar epithelium indicate the significance of the traction forces at the cell-substrate interface (Kim et al., 2013; Tambe et al., 2011; Trepat et al., 2009). However, in a pseudostratified epithelium it is unknown whether apical or basal cells generate forces needed to initiate collective migration. This knowledge gap limits our understanding of the physical mechanisms regulating the UJT.

Using pseudostratified human bronchial epithelium, we previously reported that collective cellular migration through the UJT is constrained by geometric elongation of the cells (Atia et al., 2018; Mitchel et al., 2020; O’Sullivan et al., 2020; Park et al., 2015). These studies in pseudostratified epithelia exclusively measured cell shapes at the apical surface. Due to the complexity of the physical structure of pseudostratified epithelia, intercellular forces driving apical cell elongation and collective cellular migration have not been measured. Specifically, traction forces and intercellular stresses generated by specific cell types within the pseudostratified epithelial layer are currently unmeasurable due to incompatibility of ALI culture and traction force microscopy techniques.

In our previous work using a computational model simulating two-dimensional epithelial cell boundaries, we predicted that an increase in propulsive forces induces UJT. Our computational model predicted both cell elongation and increased migration speed similar to those experimentally observed from phase-contrast microscopy of the pseudostratified epithelium (Mitchel et al., 2020). In this pseudostratified epithelium, propulsive forces can be propagated by traction forces generated at the interface of the substrate and basal cells. Thus, we hypothesize that basal cells are a source of the increased propulsive forces during the UJT. To test this hypothesis, we examined basal cell shape and actin stress fiber remodeling, which are surrogate metrics for traction force generation (Saraswathibhatla and Notbohm, 2020; Saraswathibhatla et al., 2021). Remodeling of basal cells and their stress fibers during the UJT suggests their traction force generation. In pseudostratified epithelia, basal stem cells may be responsible for initiating and propagating cellular migration during the UJT.

## Results

### During the unjamming transition, basal cells enlarge and elongate

To determine cell shape characteristics of human bronchial epithelial cells during the UJT, we measured cellular area and aspect ratio. As previously reported, we used an in vitro system of well-differentiated, pseudostratified human bronchial epithelial (HBE) cells maintained in air-liquid interface (ALI) (**Fig. 1A**) (Mitchel et al., 2016; Mitchel et al., 2020; Park and Tschumperlin, 2009; Park et al., 2015; Tschumperlin et al., 2003; Tschumperlin et al., 2004). We induced UJT by exposure to mechanical compression (**Fig. 1B**). To visualize cell morphology, we acquired z-stacks of immunofluorescence (IF) images of cells stained for F-actin. F-actin staining visualized both cell boundaries and stress fibers. While cell boundaries marked by cortical actin were prominent through all focal planes, stress fibers were only present in the regions closest to the substrate, below the nuclei of basal cells. Distinct apical cell, basal cell, and basal stress fiber regions through the apicobasally polarized epithelium were visualized using maximum intensity projections (**Fig. 1C**, detailed in *Materials and Methods*). To trace cell boundaries, we used marker-controlled watershed segmentation implemented in MATLAB (R2021a, Natick, MA).

**Figure 1.**
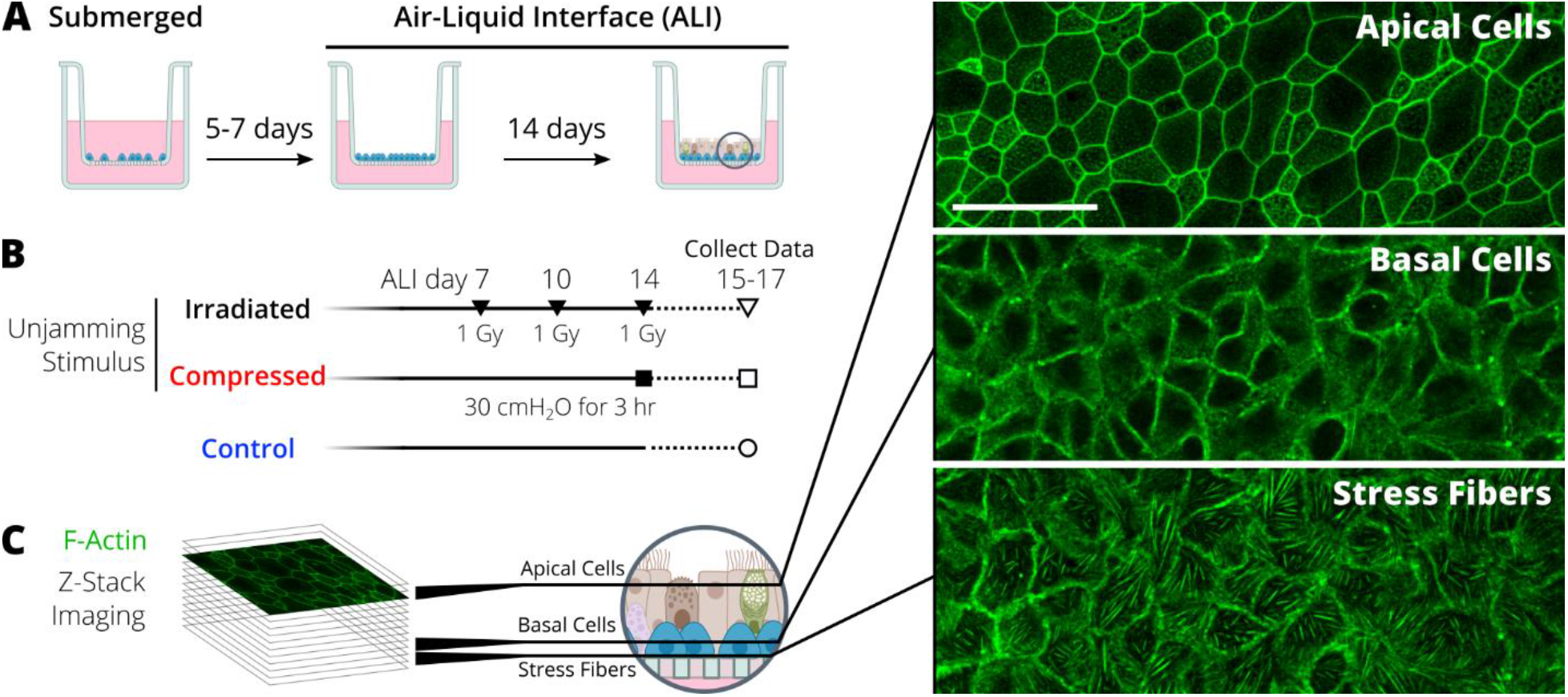
Outline of the experiment for visualizing cells and stress fibers during the unjamming transition (UJT). (A) Primary human bronchial epithelial cells were cultured in submerged and then air-liquid interface (ALI) conditions to achieve a well-differentiated, pseudostratified epithelium. (B) The layer was exposed to an unjamming stimulus (either compression or irradiation). Imaging data were collected at 24-72 hours (ALI days 15-17) after the final stimulus. (C) F-actin was stained and imaged in a z-stack. Maximum intensity projections were created to visualize different regions of interest through the apicobasally polarized epithelium. Scale bar is 50 µm.

To confirm the compressed epithelium became unjammed, we measured apical cell shape change, a hallmark of cell density-independent UJT (Atia et al., 2018; Mitchel et al., 2020; O’Sullivan et al., 2020; Park et al., 2015). From the segmented apical cell boundaries, we first quantified apical cell density (**Fig. 2A**). In the control jammed layer, cell density remained constant over 72 hours. After compression, apical cell density decreased at 48 hours but recovered by 72 hours. However, these changes in cell density showed no statistical difference between jammed (control) and unjammed (compressed) epithelium. Next, we measured apical cell area (**Fig. 2B**). In the control jammed layer, cell area remained constant through 72 hours. After compression, cell area enlarged at 48 hours but recovered to control values by 72 hours. However, these changes in cell area showed no statistical difference between jammed (control) and unjammed (compressed) epithelium. Lastly, we measured apical cell aspect ratio (AR) to characterize cell shape change as a hallmark of UJT (**Fig. 2C**). In the control jammed layer, AR remained constant over 72 hours. After compression, AR significantly increased by 48 hours (control: 1.58±0.05 vs compressed: 2.09±0.17, p<0.05). This compression-induced apical cell elongation was sustained through 72 hours. Together, we confirmed that compression-induced unjamming occurs independent of changes in apical cell density and area over 72 hours and is marked by significant apical cell elongation.

**Figure 2.**
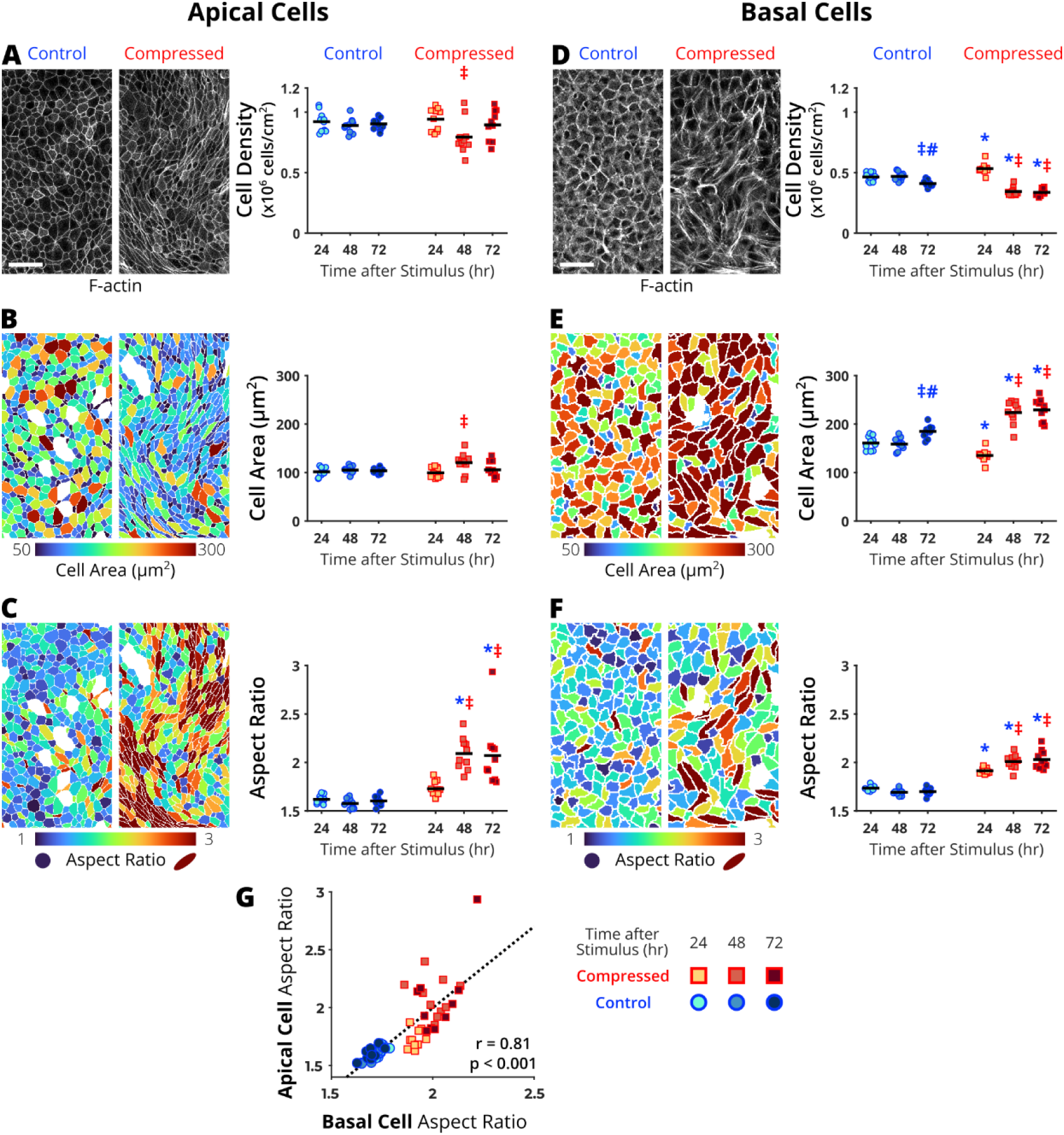
In pseudostratified airway epithelia, apical and basal cells undergo morphological changes during mechanical compression-induced unjamming transition (UJT). The representative maximum intensity projections (from cells stained for F-actin) and cell metric heatmaps are from 72 hours after control or mechanical compression. During the UJT, apical cells remained constant (A) in cell density and (B) in cell area but became (C) elongated. Basal cells (D) decreased in cell density, (E) increased in cell area, and became (F) elongated. (G) At all measured timepoints, apical and basal cell elongation were significantly correlated. Each data point represents the average value over a field of view (n=10 fields of view per time point and treatment). Significant differences are indicated for p<0.05 from one-way ANOVA with Turkey-Kramer post-hoc test (*different from time-matched control, ^‡^different from 24 hours within treatment group, ^#^different from 48 hours within treatment group). Correlations were calculated using Pearson correlation coefficient (r). Scale bar is 50 µm in (A) and (D).

Using the same IF z-stacks where we detected apical cell elongation, we characterized changes in the basal stem cells (**Fig. 1C**). In pseudostratified epithelia, cell boundaries visualized at the apical surface feature differentiated cells. However, at the basal-substrate interface, visualized cell boundaries feature mostly undifferentiated basal stem cells (**Fig. 1C**) (Evans et al., 2001; Rock et al., 2010). Despite their critical role as stem cells during development and repair, their physical characteristics during cellular migration have not been studied. We characterized morphological changes in these basal cells using the same metrics applied to the apical cells. Basal cell boundaries were visualized approximately 13±3 µm below the apical cell surface (towards the substrate, **Fig. 1C**). From the segmented basal cell boundaries, we first quantified cell density (**Fig. 2D**). In the control jammed layer, basal cell density decreased at 72 hours. In the compressed unjammed compressed layer, basal cell density first increased at 24 hours then significantly decreased at 48 and 72 hours compared to time-matched control. Next, we measured cell area (**Fig. 2E**). In the control jammed layer, basal cell area enlarged at 72 hours. In the compressed unjammed layer, basal cell area decreased at 24 hours (control: 161±14 vs compressed: 135±13 µm^2^, p<0.05). Basal cells then significantly enlarged by 48 hours (224±24 µm^2^) and plateaued through 72 hours (229±23 µm^2^). Together, these measurements indicate that basal cells are effectively decreasing their cell density by enlarging their cell area at the substrate interface during the UJT. This occurred in contrast with the apical cells which recovered cell density and area similar to control by 72 hours after compression.

Similar to the apical cells, we lastly characterized basal cell shape change by measuring AR (**Fig. 2F**). In the control jammed layer, AR remained constant over 72 hours. In the compressed unjammed layer, AR increased by 24 hours (control: 1.73±0.03 vs compressed: 1.92±0.03, p<0.05). Basal cells further elongated at 48 hours (2.01±0.07) and plateaued through 72 hours (2.03±0.09). Additionally, basal cell elongation was highly correlated with apical cell elongation (r=0.81, p<0.001, **Fig. 2G**). Together, our measurements of basal cell elongation and enlargement indicate that basal stem cells undergo remodeling during the UJT.

### Stress fibers lengthen and align during the unjamming transition

Our data show that basal stem cells in pseudostratified epithelia undergo remodeling during the UJT. These cell shape changes may play a critical role for the initiation of collective cellular migration during UJT. Therefore, to further characterize basal cell remodeling in the unjammed pseudostratified epithelium, we measured stress fiber remodeling in the basal cells. Stress fibers have been shown to be respond to mechanical changes by generating traction forces (Burridge and Guilluy, 2016; Sala and Oakes, 2021; Walcott and Sun, 2010; Zhang et al., 2017). In a simple columnar epithelium modeled by MDCK cells, unjamming-associated cell shape change and migration are controlled by stress fibers that generate traction forces at the cell-substrate interface (Saraswathibhatla and Notbohm, 2020). Furthermore, stress fiber alignment has been shown to integrate cytoskeletal forces to drive directed cell migration (Fischer et al., 2021). However, stress fiber remodeling and its link to the UJT in pseudostratified epithelia is unknown. To gain insight towards the physical forces driving migration in pseudostratified epithelia, we visualized actin stress fibers 2.2±0.4 µm below the focal plane where basal cell boundaries were visualized (**Fig. 1C**). Stress fibers exhibited a significant qualitative difference between control jammed and compressed unjammed layers (**Fig. 3A**). In the control jammed layer, stress fibers were confined within each of the cell boundaries prominently marked by cortical actin. However, in the compressed unjammed layer, stress fibers spanned multiple cell lengths and the cell boundaries were less prominent.

**Figure 3.**
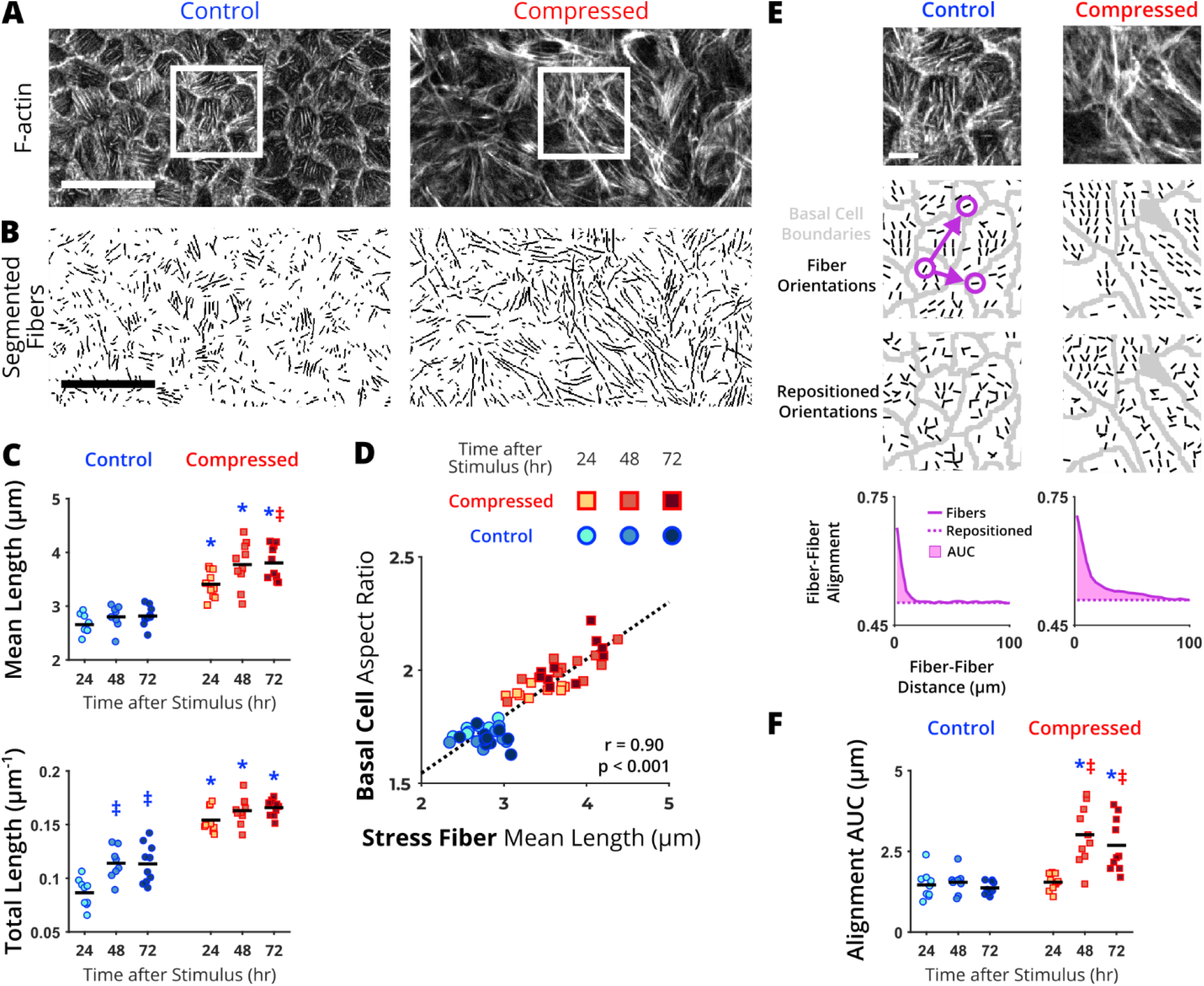
Quantification of stress fibers lengthening and alignment during the unjamming transition (UJT). (A) In a focal plane below the basal cell boundaries, we visualized stress fibers marked by F-actin staining. These representative images from control jammed and compressed unjammed epithelium were taken at 72 hours after UJT stimulus. (B) Stress fiber lengths were segmented. (C) Mean fiber lengths and total lengths all increased during the UJT.(D) Basal cell elongation and stress fiber lengthening were significantly correlated. (E) To quantify stress fiber alignment, orientation fields were determined within segmented basal cell boundaries. The regions shown in (E) are indicated the inset in (A). Fiber alignment was analyzed by plotting the average alignment between pairs of fibers against the distance between them (e.g., purple circles and arrows) for both the experimental and repositioned orientations and measuring the area under the curve (AUC, detailed in *Materials and Methods*). (F) Alignment AUC increased during the UJT. Each data point represents the average value over a field of view (n=10 fields of view per time point and treatment). Significant differences are indicated for p<0.05 from one-way ANOVA with Turkey-Kramer post-hoc test (*different from time-matched control, ^‡^different from 24 hours within treatment group, ^#^different from 48 hours within treatment group). Correlations were calculated using Pearson correlation coefficient (r). Scale bars are 50 µm in (A) and (B) and 10 µm in (E).

To characterize lengthening of stress fibers, we segmented fiber lengths using a combination of two published algorithms— FSegment (Rogge et al., 2017) and Stress Fiber Extractor (SFEX) (Zhang et al., 2017). In the control layers, we masked the visible cell boundaries to exclude potential segmentation of cortical actin and segmented solely the stress fibers (**Fig. 3B**). We calculated mean length of individual fibers and total length across each field of view. In the control jammed layer, mean stress fiber lengths remained constant over time (**Fig. 3C**). However, the total length of stress fibers increased over time (24 hours: 0.09±01 and 72 hours: 0.11±0.01 µm^-1^). In the compressed unjammed layer, mean stress fiber lengths increased at 24 hours (control: 2.7±0.2 vs compressed: 3.41±0.24 µm, p<0.05), continued to increase at 48 hours, and plateaued by 72 hours. Similarly, total stress fiber lengths increased at 24 hours (control: 0.09±0.01 µm^-1^ vs compressed: 0.15±0.01 µm^-1^, p<0.05) and continued to increase over 72 hours. Together, these data indicate that stress fibers are significantly lengthening and accumulating during the UJT. Interestingly, the time course of stress fiber lengthening was significantly correlated with basal cell elongation (r=0.90, p<0.001), suggesting a causal relationship between stress fiber and basal cell remodeling during the UJT (**Fig. 3D**).

To characterize organization of stress fibers, we used the software MatFiber to estimate stress fiber orientations within basal cells (**Fig. 3E**, detailed in *Materials and Methods*) (Fomovsky and Holmes, 2010). The orientation vector fields were used to quantify regional heterogeneity of alignment (Richardson and Holmes, 2016). In the control jammed layer, the stress fibers maintain their heterogeneous alignment measured using alignment area under the curve (AUC) (**Fig. 3F**). In the compressed unjammed layer, alignment AUC significantly increased by 48 hours (control: 1.5±0.4 vs compressed: 3.0±0.9 µm^-1^, p<0.05). The increased alignment was sustained through 72 hours of the UJT. Together, these results demonstrate that during compression-induced UJT, basal cell stress fibers lengthen by 24 hours and align by 48 hours. This stress fiber remodeling suggests that basal cells are a source of traction forces during compression-induced UJT.

### Basal cell and stress fiber changes are novel hallmarks of the unjamming transition

With our data revealing significant correlations between apical cell elongation, basal cell remodeling, and stress fiber remodeling in pseudostratified epithelia, we further hypothesized that they are common characteristics of the UJT regardless of the stimuli. To test this hypothesis, we performed a study using either mechanical compression or ionizing radiation to induce UJT (**Fig. 1B**) (Mitchel et al., 2020; O’Sullivan et al., 2020; Park et al., 2015). After inducing UJT, we first measured both dynamic and structural characteristics of the UJT. The dynamic characteristics of the UJT includes cell migration. As in previous work, both UJT stimuli induced collective cellular migration after 72 hours as visualized by spatially coordinated migration paths and speeds (**Fig. 4A**). In the control jammed layer, the migration paths over a 1.5-hour period remained centered on the initial tracking grid. After induction of UJT by either compression or irradiation, the migration paths exhibited dramatic swirling patterns with distinct clusters of high and low speeds indicative of cellular unjamming (Angelini et al., 2011; Mitchel et al., 2020; Palamidessi et al., 2019; Park et al., 2015; Vishwakarma et al., 2020). In the control jammed layer, average migration speeds were low (24 hours: 0.13±0.03 and 72 hours: 0.23±0.04 µm/hr, **Fig. 4B**). After both compression and irradiation, migration speeds significantly increased by 72 hours (compressed: 1.77±1.25 µm/hr; irradiated: 4.10±1.42 µm/hr).

**Figure 4.**
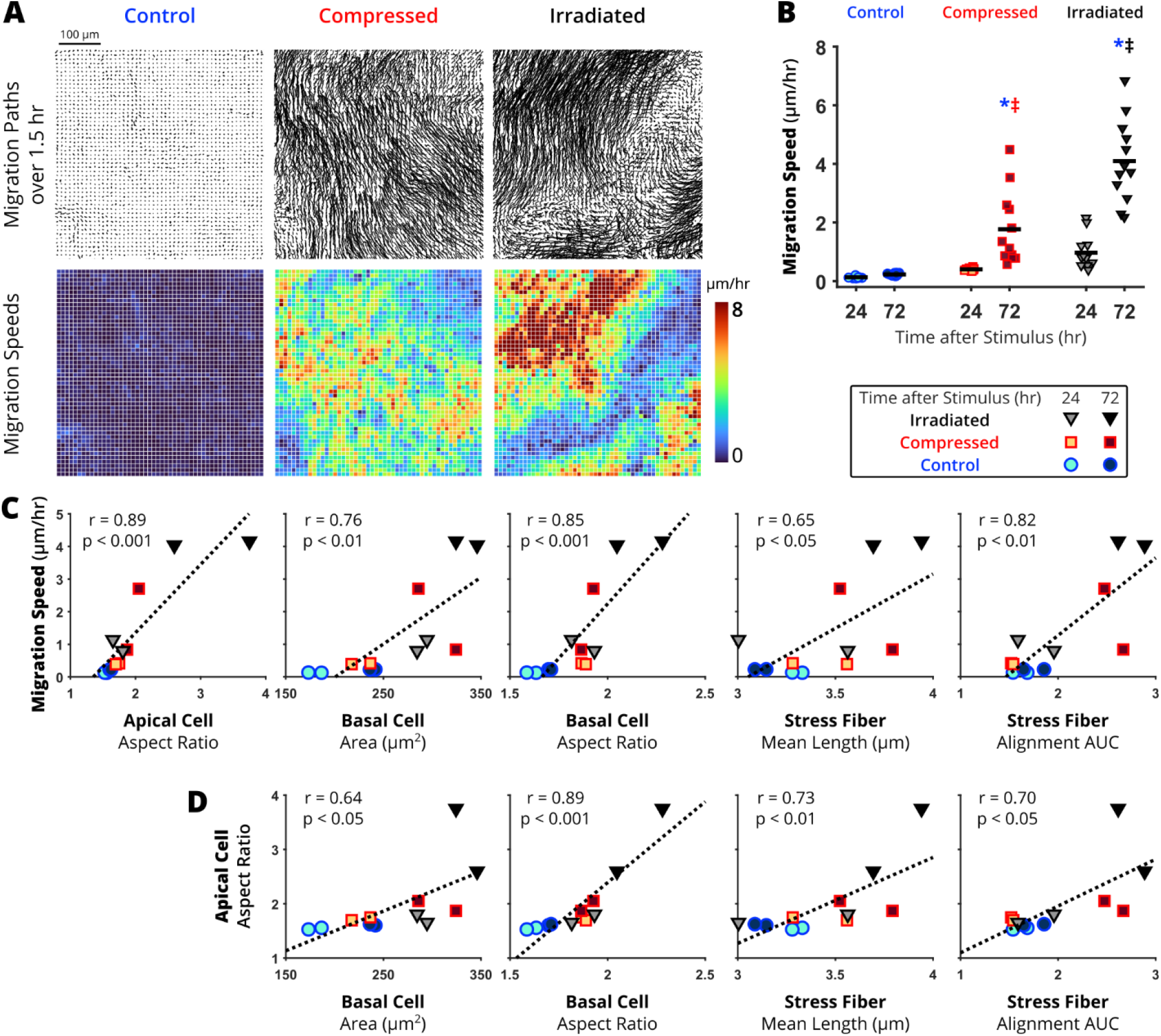
Basal cell remodeling metrics correlate with migration speed and apical cell elongation during the unjamming transition. Unjamming was induced by either mechanical compression or irradiation. At 24 or 72 hours after exposure to either stimulus, migration speed was measured from time-lapse phase microscopy images. (A) In representative fields of view at 72 hours, migration paths were traced to calculate speeds starting from a 10 µm grid over 1.5 hours. (B) At 24 and 72 hours, the average migration speed over each field of view was calculated for the jammed (control) and unjammed (compressed and irradiated) layers.Significant differences are indicated for p<0.05 from one-way ANOVA with Turkey-Kramer post-hoc test (*different from time-matched control, ^‡^different from 24 hour within treatment group, ^#^different from 48 hour within treatment group). (C) Migration speeds were correlated with apical cell elongation, basal cell enlargement and elongation, as well as stress fiber lengthening and alignment. (D) Apical cell elongation was correlated with basal cell enlargement and elongation as well as stress fiber lengthening and alignment. Each data point in (C-D) represents the average across one experimental well. Correlations were calculated using Pearson correlation coefficient (r) with n=2 wells per treatment and timepoint.

We quantified the correlation between our newly characterized basal cell and stress fiber morphology metrics and migration speed and apical cell aspect ratio, two previously published hallmarks of the UJT (Mitchel et al., 2020; Park et al., 2015). During the UJT, migration speed was significantly correlated with apical cell elongation (r=0.89, p<0.001), basal cell enlargement (r=0.76, p<0.01), basal cell elongation (r=0.85, p<0.001), stress fiber lengthening (r=0.65, p<0.05), and stress fiber alignment (r=0.82, p<0.01) (**Fig. 4C**). Additionally, apical cell elongation was correlated with basal cell enlargement (r=0.64, p<0.05), basal cell elongation (r=0.89, p<0.001), stress fiber lengthening (r=0.73, p<0.01), and stress fiber alignment (r=0.70, p<0.05) (**Fig. 4D**). The significant correlation of basal cell and stress fiber metrics with migration speed and apical cell elongation for two, independent UJT stimuli support basal cell and stress fiber remodeling as novel structural hallmarks of the UJT in pseudostratified epithelia.

## Discussion

Collective cellular migration is a critical cellular process in development and disease (Crosby and Waters, 2010; Goodwin and Nelson, 2021; Varner and Nelson, 2014). One mechanism for epithelial cell migration is through the unjamming transition (UJT) (Atia et al., 2021; Buttenschön and Edelstein-Keshet, 2020; Garcia et al., 2015; Kim et al., 2020; La Porta and Zapperi, 2020; Oswald et al., 2017; Palamidessi et al., 2019; Park et al., 2016). UJT in the pseudostratified airway epithelium has previously been characterized by apical cell elongation and collective cellular migration (Atia et al., 2018; Mitchel et al., 2020; Park et al., 2015). While these characteristics of the UJT suggest the presence of intercellular force changes, the physical forces driving collective cellular migration in the UJT are unknown. Despite the heterogeneity of pseudostratified airway epithelium composed of both differentiated apical cells and undifferentiated basal stem cells, the cell-type specific characteristics during UJT have not been studied. In pseudostratified airway epithelia, multiple types of differentiated epithelial cells are in contact with the basal lamina (or substrate *in vitro*); however, basal stem cells occupy most of the basal lamina area (Evans et al., 2001). Because of this prominent contact, we hypothesized that basal cells generate tractions forces necessary for migration during the UJT. To interrogate the role of basal cells in the UJT, we used image analysis to characterize morphological changes in basal cells and their stress fibers. Our results reveal that basal stem cells actively remodel as part of the UJT in pseudostratified airway epithelium.

We first induced UJT in an *in vitro* model of airway epithelium by applying mechanical compression. Our data unveil four novel hallmarks of the UJT— basal cell enlargement and elongation as well as stress fiber lengthening and alignment. By 24 hours of UJT, basal cells elongated (**Fig. 2E**) and their stress fibers lengthened (**Fig. 3C**). By 48 hours, basal cells enlarged (**Fig. 2F**) and their stress fiber aligned (**Fig. 3F**). Together, our data demonstrate that basal cells and their stress fibers actively remodel during the UJT. Importantly, basal cell remodeling preceded apical cell elongation (at 48 hours, **Fig. 2B**), a previously established hallmark of the UJT (Atia et al., 2018; Mitchel et al., 2020; Park et al., 2015). Interestingly, our previous studies showed that apical cell elongation significantly increased by 24 hours after compression. The delay in significant apical cell elongation in the current data set may be due to donor-to-donor variability when using primary HBE cells.

We validated the four novel hallmarks of the UJT in an additional dataset collected using either compression- or irradiation-induced UJT (**Fig. 4**). Mechanical compression and ionizing radiation induced UJT with different magnitudes of migration speed and apical cell elongation (**Fig. 4B**). And the basal cell and stress fiber metrics were all significantly correlated with migration speed (**Fig. 4C**) and apical cell elongation (**Fig. 4D**), two previously established hallmarks of UJT (Mitchel et al., 2020; O’Sullivan et al., 2020; Park et al., 2015). These novel hallmarks expand the current definition and interpretation of UJT in pseudostratified epithelia. Our prior studies have focused on the role of cell shape changes at the apical surface. Here our current study demonstrates that even before the onset of cellular migration or apical cell elongation, basal cells are actively remodeling during the UJT in pseudostratified airway epithelium. Therefore, how basal cells undergo remodeling and how they play a role during UJT should be further investigated to advance our mechanistic insight toward the UJT.

During the UJT, apical cells maintained constant cell area and density (**Fig. 2A and 2C**). However, basal cells enlarged and decreased their density (**Fig. 2D and 2F**). We have also previously shown that during the UJT epithelial barrier integrity remains intact (Mitchel et al., 2020). Together, our data reflect previously reported observations of basal cell remodeling in pseudostratified airway epithelium in airway diseases or after injury. For example, airway basal cells flatten (or increase cell area) in response to damage and loss of differentiated cells above them to preserve barrier integrity (Erjefält et al., 1997; Puchelle et al., 2006). Accompanying the basal cell enlargement, our data indicate that basal cell density decreases during UJT; however, the mechanism for basal cell loss remains unclear. We speculate that basal cells may have been extruded during UJT. Or these basal stem cells may be differentiating to replace any loss of apical cells (through cell death or extrusion) (Baba et al., 2020; Gudipaty and Rosenblatt, 2017; Kretschmer et al., 2017; Tadokoro et al., 2016). In monolayer cell culture, increasing cell density has been shown to decrease the speed of collective cellular migration (Angelini et al., 2011; Garcia et al., 2015; Loza et al., 2016; Saraswathibhatla and Notbohm, 2020; Tambe et al., 2011; Tlili et al., 2018). However, in pseudostratified epithelium, changes in density were not previously explored or characterized during the UJT. Thus, our data demonstrating changes in basal cell density raise new questions for the UJT.

The physical forces driving UJT in pseudostratified epithelium are not well understood due to limitations in methods to directly measure traction forces in ALI culture. Prior computational modeling studies have suggested that increased cell-cell adhesion at the cell periphery can induce the UJT (Bi et al., 2014; Park et al., 2015; Park et al., 2016). In the case of pseudostratified epithelium, UJT has been attributed to changes in cortical tension or adhesions at the cell boundaries; however, the observations supporting this mechanism have been measured exclusively at the apical surface (Mitchel et al., 2020; O’Sullivan et al., 2020; Park et al., 2015). Computational modeling has also suggested that increasing cellular force generation (propulsive forces) in the model also induces the UJT in the absence of changes to cortical tension or adhesion (Mitchel et al., 2020). Experimentally, traction forces have been directly measured in a monolayer of cells and shown to be critical for cell shape change and migration during UJT (Saraswathibhatla and Notbohm, 2020). Specifically, traction forces are generated by actin stress fibers and lead to cell elongation suggesting the significance of forces at the cell-substrate interface. In the absence of traction force measurements, cell shape elongation and stress fiber accumulation and alignment are reliable surrogate metrics for traction force generation (Saraswathibhatla and Notbohm, 2020; Vignaud et al., 2020). Our data indicate that in pseudostratified epithelia, basal cells and their stress fibers at the substrate interface undergo remodeling suggestive of traction force generation during the UJT (**Fig. 5**). These cellular forces induced by accumulated stress fibers may be the driving mechanism for the previously observed apical cell elongation and collective cellular migration. However, the physical mechanism connecting basal cell traction forces with apical cell elongation remains unclear.

**Figure 5.**
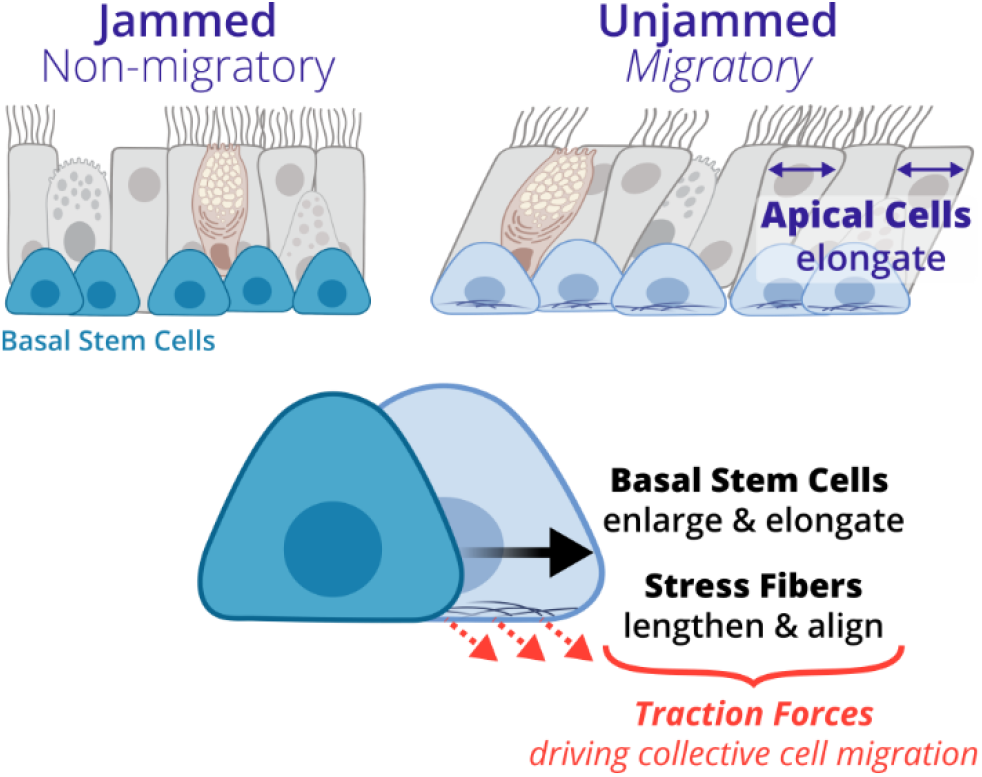
A conceptual schematic of the unjamming transition (UJT) in pseudostratified airway epithelium. UJT has been previously characterized by increased collective cellular migration and apical cell elongation (Park et al., 2015). In this study, we now identified that basal stem cells undergo unique structural changes, including cell elongation and enlargement as well as stress fiber lengthening and alignment during the UJT. These findings suggest basal stem cells are generating traction forces required for driving collective cellular migration during the UJT.

Apical cells in a pseudostratified epithelium are tethered together via a combination of tight junctions and adherens junctions, and basal cells are tethered to the apical cells via desmosomes (Evans et al., 2001; Niessen, 2007; Roche et al., 1993; Rock et al., 2010). As the basal cells generate traction forces, desmosomes may be relaying these forces to the apical cells to induce the previously described hallmarks of the UJT (Mitchel et al., 2020; O’Sullivan et al., 2020; Park et al., 2015). Desmosome assembly in a migrating cell monolayer has been shown to localize at the leading edge of migration and require actin cytoskeleton force generation (Koetsier et al., 2014; Roberts et al., 2011; Thomason et al., 2010). Additionally, the role of cell neighbors throughout the apicobasal axis of the epithelium may be critical to understanding intercellular force transmission and unjamming (Gómez et al., 2021; Grosser et al., 2021). The role of desmosomes in physical tethering of basal and apical cells as well as the three-dimensional organization of cells should be further studied in unjammed pseudostratified epithelium.

In this study, our novel morphological analyses of basal stem cells and actin stress fibers reveal that basal cells actively remodel and may generate traction forces driving collective cellular migration during the UJT. We identified new hallmarks of the UJT: basal cell enlargement and elongation as well as stress fiber lengthening and alignment. These hallmarks expand the library of metrics that can be used to characterize the jamming or unjamming of pseudostratified epithelium in both *in vitro* and *in vivo* models. Furthermore, this work supports future mechanistic studies towards the intracellular signaling by which basal stem cells regulate traction force generation. While basal stem cells have been well-recognized as a rich source of biochemical signals as epithelial progenitor cells, our new findings recognize a critical, physical role for basal stem cells as an active driver of collective cellular migration.

## Materials and methods

### Culture of primary human bronchial epithelial cells

Primary human bronchial epithelial (HBE) cells were obtained from the Marisco Lung Institute/Cystic Fibrosis Research Center at the University of North Carolina at Chapel Hill (UNC) under protocol No. 03-1396 approved by the UNC Biomedical Institutional Review Board. The two donors used in this study were from non-smokers with no history of chronic lung disease. As described previously (Mitchel et al., 2016; Mitchel et al., 2020; O’Sullivan et al., 2020; O’Sullivan et al., 2021; Park and Tschumperlin, 2009; Park et al., 2010; Park et al., 2012; Park et al., 2015), HBE cells at passage 2 were cultured on transwell inserts (Corning, 12 mm, 0.4 µm pore, polyester) and maintained in air-liquid interface (ALI) for 14 days until well-differentiated (**Fig. 1**).

To induce the unjamming transition (UJT), we exposed cells to either mechanical compression or ionizing radiation (Figure 1) (Mitchel et al., 2020; O’Sullivan et al., 2020; Park et al., 2015). For mechanical compression, cells were exposed to an apical-to-basal pressure differential of 30 cmH_2_O for 3 hours on ALI day 14 (Mitchel et al., 2016; Mitchel et al., 2020; Park and Tschumperlin, 2009; Park et al., 2010; Park et al., 2012; Park et al., 2015; Tschumperlin et al., 2004). For irradiation, cells were exposed to a dose of 1 Gy of ionizing radiation using a RS 2000 Biological Research Irradiator (RadSource, Brentwood, TN) on ALI culture days 7, 10, and 14 (O’Sullivan et al., 2020). Following the final unjamming stimulus on ALI day 14, cells were maintained in ALI culture for up to 72 hours.

In the cells from donor one, we applied mechanical compression to induce the UJT and collected imaging data after 24, 48, or 72 hours. In the cells from donor two, we induced unjamming using either mechanical compression or irradiation and collected imaging data 24 or 72 hours after each of the unjamming stimuli. For all experiments, two transwells were imaged for each treatment and timepoint.

### Immunofluorescence staining and static imaging

At 24, 48, or 72 hours after final exposure to the unjamming stimulus, cells were fixed with 4% paraformaldehyde in PBS with calcium and magnesium for 30 minutes at room temperature. Cells were permeabilized with 0.2% Triton X-100 for 15 minutes, blocked with 1% bovine serum albumin and 10% normal goat serum for 1 hour, and stained for F-actin (Alexa fluor 488-Phalloidin, ThermoFisher Scientific, diluted 1:40, 30 minutes). Transwell membranes were cut from the plastic support and mounted on glass slides (Vectashield). Slides were imaged using a Zeiss Axio Observer Z1 with an apotome module controlled using Zen Blue 2.0 software. Five random fields of view (563 by 356 µm) were imaged from each transwell membrane in a z-stack from substrate to apical cell surface (15±3 µm in height).

To visualize various planes through the pseudostratified epithelial layer (**Fig. 1**), maximum intensity projections were generated in Fiji (Schindelin et al., 2012). The apical surface was generated from all z-stack slices that included visible apical boundaries. Approximately 13±3 µm below the apical surface, the basal cell boundaries were manually masked using a sliding region of interest across the field of view in sequential z-stack slices before a maximum intensity projection was generated. To visualize stress fibers, the identical regions of interest for the basal cells were masked in lower z-stack slices (2.2±0.4 µm) towards the transwell membrane before generating a maximum intensity projection.

### Cell shape analysis

Cell boundaries were visualized from maximum intensity projections of apical and basal surfaces, generated as described above, using marker-controlled watershed segmentation in MATLAB (R2021a). Apical cell images were first filtered using opening-closing by reconstruction. After initial watershed segmentation, quality control of the segmented cell borders was assessed by calculating boundary tortuosity, length, and mean pixel intensity. Over-segmented boundaries were removed based on thresholds of the three metrics. Similarly, basal cell images were first filtered using a Wiener filter followed by watershed segmentation. Apical and basal cell bodies were lastly filtered based on area to remove under- and over-segmented regions.

Cell morphology was characterized by measuring cell area and aspect ratio. Cell area was quantified in the plane parallel to the transwell membrane (Mitchel et al., 2020; O’Sullivan et al., 2020; Park et al., 2015). Cell aspect ratio was defined as the ratio of the major- to minor-axis length for an ellipse fit to the segmented cell boundary. These metrics were mapped to the segmentation for visualization of morphology changes. For comparison between treatments and timepoints, cell morphology metrics were averaged over each field of view.

### Stress fiber analysis

Stress fibers lengths were segmented using a combination of two published algorithms— FSegment (Rogge et al., 2017) and Stress Fiber Extractor (SFEX) (Zhang et al., 2017). Briefly, the FSegment algorithm is designed to iteratively trace fiber fragments within an image (Rogge et al., 2017). The iterative design specifically overcomes the challenge of superimposed stress fibers. FSegment outputs fiber fragments which often separate longer fibers into multiple shorter regions. We used the fiber reconstruction algorithm developed in SFEX to connect the fragments.Briefly, SFEX evaluates the end points of all fiber fragments and connects them if they meet geometric constraints (based on distance and orientation of fiber fragments) (Zhang et al., 2017). Additionally, we added a constraint to check whether potential connecting segments between fiber fragments featured pixel intensities matching the fiber fragments. In the control groups, cortical actin at the cell boundaries confounded the stress fiber segmentation (**Fig. 3A**); therefore, we removed the basal boundary regions segmented from the basal cell plane image before segmenting fibers. In the unjammed groups, cortical actin was not prevalent; therefore, we did not mask the basal cell boundaries. To measure the mean length of stress fibers, we averaged the lengths across a field of view. To measure the total length of stress fibers, we summed the lengths and divided by the size of the field of view searched by the segmentation algorithm.

To analyze alignment of stress fibers, we quantified fiber orientations using the software MatFiber and measured spatial heterogeneity (Fomovsky and Holmes, 2010; Richardson and Holmes, 2016). To isolate the stress fibers from the cell boundaries, we removed the basal cell boundaries segmented from the basal plane image and dilated to a width of 1.5 µm (**Fig. 3E**). We evaluated the orientation of stress fibers in finite subregions (3 by 3 µm) of the image with MatFiber. MatFiber uses an intensity-gradient-detection algorithm to measure orientation of fibers. To exclude subregions without stress fibers, we removed subregions with mean pixel intensity below the 35^th^ percentile based on the basal cell boundary-free image. To evaluate the spatial heterogeneity of stress fiber orientations, we calculated the Alignment Area Under the Curve (AUC) metric described in detail by (Richardson and Holmes, 2016). From the experimental orientation field, we plotted the average alignment (dot product) between pairs of fibers against the distance between them. We then randomly repositioned all the orientation vectors to remove local alignment and repeated the plotting. The area captured between the two curves (Alignment AUC) is a single measure that characterizes the degree to which local alignment of fibers exceeds global alignment across the image. Higher values of Alignment AUC indicate that stress fibers are more similarly aligned locally.

### Live imaging and cellular migration analysis

Cellular migration speeds were measured from time-lapse images taken at 24 or 72 hours after unjamming stimulus treatment. For each independent experimental replicate (2 transwells per treatment per timepoint), six fields of view (1124 by 713 µm) per well were imaged every 6 minutes over 1.5 hours. The imaging chamber was supplied with 37°C, 5% CO_2_, humidified air on a Zeiss Axio Observer Z1 to collect phase contrast images. Flow fields were calculated using optical flow with the Farneback method in MATLAB (Farnebäck, 2003). A 10 by 10 µm grid was initially seeded in the first image, and the migration trajectories were calculated by forwards-integration of the flow field. The average migration speed was calculated from the displacement over the 1.5-hour time-lapse.

### Statistics

For each treatment and timepoint, two wells were imaged with five fields of view for z-stacks and six fields of view for time-lapses. All statistics and visualizations were computed in MATLAB. To compare metrics across treatments and timepoints, we averaged the metric across each field of view and used a one-way ANOVA with Tukey-Kramer post-hoc test. Groups were considered statistically significant for p<0.05. For data visualization, we show a data point for each field of view with a solid line in each group to indicate the mean value. In the text, metrics are reported as mean ± standard deviation.

For correlations between metrics, we calculated a Pearson correlation coefficient (*r*) and report the p-value. The correlations between cell morphology and stress fiber metrics (**Fig. 2G and 3D**) used data points from matched z-stacks. The correlations with migration speeds (**Fig. 4C**) or apical cell aspect ratio (**Fig. 4D**) used data points averaged across each transwell. To get a single value for each transwell, the cell and stress fiber morphology metrics were averaged across five fields of view per transwell and the migration speeds were averaged across six fields of view per transwell. For data visualization, we show the individual data points with a dashed line indicating the line of best fit.

## Acknowledgements

Schematics in **Figs. 1 and 5** were created with Biorender.com.

## Data availability

The imaging datasets are available on Zenodo: **Figs. 2 and 3** (DOI: 10.5281/zenodo.7005090), **Fig. 4** (DOI: 10.5281/zenodo.7005123). Image analysis code is available on Zenodo: cell segmentation (DOI: 10.5281/zenodo.6998356), stress fiber segmentation (DOI: 10.5281/zenodo.6998358), stress fiber alignment heterogeneity (DOI: 10.5281/zenodo.6998361), and cell migration speed (DOI: 10.5281/zenodo.6999304).

## References

Angelini, T. E., Hannezo, E., Trepatc, X., Marquez, M., Fredberg, J. J. and Weitz, D. A. (2011). Glass-like dynamics of collective cell migration. Proc. Natl. Acad. Sci. U. S. A. 108, 4714–4719.

Atia, L., Bi, D., Sharma, Y., Mitchel, J. A., Gweon, B., Koehler, S. A., Decamp, S. J., Lan, B., Kim, J. H., Hirsch, R., et al. (2018). Geometric constraints during epithelial jamming. Nat. Phys. 14, 613–620.

Atia, L., Fredberg, J. J., Gov, N. S. and Pegoraro, A. F. (2021). Are cell jamming and unjamming essential in tissue development? Cells Dev. 168, 203727.

Baba, K., Sasaki, K., Morita, M., Tanaka, T., Teranishi, Y., Ogasawara, T., Oie, Y., Kusumi, I., Inoie, M., Hata, K., et al. (2020). Cell jamming, stratification and p63 expression in cultivated human corneal epithelial cell sheets. Sci. Rep. 10, 9282.

Bi, D., Lopez, J. H., Schwarz, J. M. and Lisa Manning, M. (2014). Energy barriers and cell migration in densely packed tissues. Soft Matter 10, 1885–1890.

Burridge, K. and Guilluy, C. (2016). Focal adhesions, stress fibers and mechanical tension Exp. Cell Res. 343, 14–20.

Butler, J. P., Toli-Nørrelykke, I. M., Fabry, B. and Fredberg, J. J. (2002). Traction fields, moments, and strain energy that cells exert on their surroundings. Am. J. Physiol. - Cell Physiol. 282, 595–605.

Buttenschön, A. and Edelstein-Keshet, L. (2020). Bridging from single to collective cell migration: A review of models and links to experiments.

Crosby, L. M. and Waters, C. M. (2010). Epithelial repair mechanisms in the lung. Am. J. Physiol. - Lung Cell. Mol. Physiol. 298,.

DeCamp, S. J., Tsuda, V. M. K., Ferruzzi, J., Koehler, S. A., Giblin, J. T., Roblyer, D., Zaman, M. H., Weiss, S. T., Kılıç, A., De Marzio, M., et al. (2020). Epithelial layer unjamming shifts energy metabolism toward glycolysis. Sci. Rep. 10, 1–15.

Erjefält, J. S., Sundler, F. and Persson, C. G. A. (1997). Epithelial barrier formation by airway basal cells. Thorax 52, 213–217.

Evans, M. J., Van Winkle, L. S., Fanucchi, M. V. and Plopper, C. G. (2001). Cellular and molecular characteristics of basal cells in airway epithelium. Exp. Lung Res. 27, 401–415.

Farnebäck, G. (2003). Two-Frame Motion Estimation Based on Polynomial Expansion. In Proceedings of the 13th Scandinavian Conference on Image Analysis, pp. 363–370.

Fischer, R. S., Sun, X., Baird, M. A., Hourwitz, M. J., Seo, B. R., Pasapera, A. M., Mehta, S. B., Losert, W., Fischbach, C., Fourkas, J. T., et al. (2021). Contractility, focal adhesion orientation, and stress fiber orientation drive cancer cell polarity and migration along wavy ECM substrates. Proc. Natl. Acad. Sci. U. S. A. 118,.

Fomovsky, G. M. and Holmes, J. W. (2010). Evolution of scar structure, mechanics, and ventricular function after myocardial infarction in the rat. Am. J. Physiol. Heart Circ. Physiol. 298, H221–8.

Garcia, S., Hannezo, E., Elgeti, J., Joanny, J. F., Silberzan, P. and Gov, N. S. (2015). Physics of active jamming during collective cellular motion in a monolayer. Proc. Natl. Acad. Sci. U. S. A. 112, 15314–15319.

Gómez, H. F., Dumond, M. S., Hodel, L., Vetter, R. and Iber, D. (2021). 3D cell neighbour dynamics in growing pseudostratified epithelia. Elife 10, 1–25.

Goodwin, K. and Nelson, C. M. (2021). Mechanics of Development. Dev. Cell 56, 240–250.

Gray, T. E., Guzman, K., Davis, C. W., Abdullah, L. H. and Nettesheim, P. (1996). Mucociliary differentiation of serially passaged normal human tracheobronchial epithelial cells. Am. J. Respir. Cell Mol. Biol. 14, 104–12.

Grosser, S., Lippoldt, J., Oswald, L., Merkel, M., Sussman, D. M., Renner, F., Gottheil, P., Morawetz, E. W., Fuhs, T., Xie, X., et al. (2021). Cell and Nucleus Shape as an Indicator of Tissue Fluidity in Carcinoma. Phys. Rev. X 11, 011033.

Gudipaty, S. A. and Rosenblatt, J. (2017). Epithelial cell extrusion: Pathways and pathologies.emin. Cell Dev. Biol. 67, 132–140.

Kim, J. H., Serra-Picamal, X., Tambe, D. T., Zhou, E. H., Park, C. Y., Sadati, M., Park, J. A., Krishnan, R., Gweon, B., Millet, E., et al. (2013). Propulsion and navigation within the advancing monolayer sheet. Nat. Mater. 12, 856–863.

Kim, J. H., Pegoraro, A. F., Das, A., Koehler, S. A., Ujwary, S. A., Lan, B., Mitchel, J. A., Atia, L., He, S., Wang, K., et al. (2020). Unjamming and collective migration in MCF10A breast cancer cell lines. Biochem. Biophys. Res. Commun. 521, 706–715.

Koetsier, J. L., Amargo, E. V., Todorović, V., Green, K. J. and Godsel, L. M. (2014).Plakophilin 2 affects cell migration by modulating focal adhesion dynamics and integrin protein expression. J. Invest. Dermatol. 134, 112–122.

Kretschmer, S., Pieper, M., Klinger, A., Hüttmann, G. and König, P. (2017). Imaging of Wound Closure of Small Epithelial Lesions in the Mouse Trachea. Am. J. Pathol. 187, 2451–2460.

La Porta, C. A. M. and Zapperi, S. (2020). Phase transitions in cell migration. Nat. Rev. Phys.2, 516–517.

Loza, A. J., Koride, S., Schimizzi, G. V., Li, B., Sun, S. X. and Longmore, G. D. (2016). Cell density and actomyosin contractility control the organization of migrating collectives within an epithelium. Mol. Biol. Cell 27, 3459–3470.

Mitchel, J. A., Antoniak, S., Lee, J. H., Kim, S. H., McGill, M., Kasahara, D. I., Randell, S. H., Israel, E., Shore, S. A., Mackman, N., et al. (2016). IL-13 augments compressive stress-induced tissue factor expression in human airway epithelial cells. Am. J. Respir. Cell Mol.Biol. 54, 524–531.

Mitchel, J. A., Das, A., O’Sullivan, M. J., Stancil, I. T., DeCamp, S. J., Koehler, S., Ocaña, O. H., Butler, J. P., Fredberg, J. J., Nieto, M. A., et al. (2020). In primary airway epithelial cells, the unjamming transition is distinct from the epithelial-to-mesenchymal transition. Nat. Commun. 11, 5053.

Montoro, D. T., Haber, A. L., Rood, J. E., Regev, A. and Rajagopal, J. (2020). A Synthesis Concerning Conservation and Divergence of Cell Types across Epithelia. Cold Spring Harb. Perspect. Biol. 12, 1–14.

Niessen, C. M. (2007). Tight junctions/adherens junctions: Basic structure and function. J. Invest. Dermatol. 127, 2525–2532.

O’Sullivan, M. J., Mitchel, J. A., Das, A., Koehler, S., Levine, H., Bi, D., Nagel, Z. D. and Park, J.-A. (2020). Irradiation Induces Epithelial Cell Unjamming. Front. Cell Dev. Biol. 8,.

O’Sullivan, M. J., Mitchel, J. A., Mwase, C., McGill, M., Kanki, P. and Park, J. A. (2021). In well-differentiated primary human bronchial epithelial cells, TGF-b1 and TGF-β2 induce expression of furin. Am. J. Physiol. - Lung Cell. Mol. Physiol. 320, L246–L253.

Oswald, L., Grosser, S., Smith, D. M. and Käs, J. A. (2017). Jamming transitions in cancer. J. Phys. D. Appl. Phys. 50,.

Palamidessi, A., Malinverno, C., Frittoli, E., Corallino, S., Barbieri, E., Sigismund, S., Beznoussenko, G. V, Martini, E., Garre, M., Ferrara, I., et al. (2019). Unjamming overcomes kinetic and proliferation arrest in terminally differentiated cells and promotes collective motility of carcinoma. Nat. Mater. 18, 1252–1263.

Park, J. A. and Tschumperlin, D. J. (2009). Chronic intermittent mechanical stress increases MUC5AC protein expression. Am. J. Respir. Cell Mol. Biol. 41, 459–466.

Park, J. A., Drazen, J. M. and Tschumperlin, D. J. (2010). The chitinase-like protein YKL-40 is secreted by airway epithelial cells at base line and in response to compressive mechanical stress. J. Biol. Chem. 285, 29817–29825.

Park, J.-A., Sharif, A. S., Tschumperlin, D. J., Lau, L., Limbrey, R., Howarth, P. and Drazen, J. M. (2012). Tissue factor–bearing exosome secretion from human mechanically stimulated bronchial epithelial cells in vitro and in vivo. J. Allergy Clin. Immunol. 130, 1375–1383.

Park, J.-A., Kim, J. H., Bi, D., Mitchel, J. A., Qazvini, N. T., Tantisira, K., Park, C. Y., McGill, M., Kim, S.-H., Gweon, B., et al. (2015). Unjamming and cell shape in the asthmatic airway epithelium. Nat. Mater. 14, 1040–8.

Park, J. A., Atia, L., Mitchel, J. A., Fredberg, J. J. and Butler, J. P. (2016). Collective migration and cell jamming in asthma, cancer and development. J. Cell Sci. 129, 3375–3383.

Puchelle, E., Zahm, J. M., Tournier, J. M. and Coraux, C. (2006). Airway epithelial repair, regeneration, and remodeling after injury in chronic obstructive pulmonary disease. Proc. Am. Thorac. Soc. 3, 726–733.

Richardson, W. J. and Holmes, J. W. (2016). Emergence of Collagen Orientation Heterogeneity in Healing Infarcts and an Agent-Based Model. Biophys. J. 110, 2266–2277.

Roberts, B. J., Pashaj, A., Johnson, K. R. and Wahl, J. K. (2011). Desmosome dynamics in migrating epithelial cells requires the actin cytoskeleton. Exp. Cell Res. 317, 2814–2822.

Roche, W. R., Montefort, S., Baker, J. and Holcate, S. T. (1993). Cell Adhesion Molecules and the Bronchial Epithelium. Am. Rev. Respir. Dis. 148, S79–S82.

Rock, J. R., Randell, S. H. and Hogan, B. L. M. (2010). Airway basal stem cells: A perspective on their roles in epithelial homeostasis and remodeling. DMM Dis. Model. Mech. 3, 545–556.

Rogge, H., Artelt, N., Endlich, N. and Endlich, K. (2017). Automated segmentation and quantification of actin stress fibres undergoing experimentally induced changes. J. Microsc. 268, 129–140.

Sala, S. and Oakes, P. W. (2021). Stress fiber strain recognition by the LIM protein testin is cryptic and mediated by RhoA. Mol. Biol. Cell 32, 1758–1771.

Saraswathibhatla, A. and Notbohm, J. (2020). Tractions and Stress Fibers Control Cell Shape and Rearrangements in Collective Cell Migration. Phys. Rev. X 10, 11016.

Saraswathibhatla, A., Galles, E. E. and Notbohm, J. (2020). Spatiotemporal force and motion in collective cell migration. Sci. Data 7, 1–7.

Saraswathibhatla, A., Henkes, S., Galles, E. E., Sknepnek, R. and Notbohm, J. (2021).Coordinated tractions increase the size of a collectively moving pack in a cell monolayer.Extrem. Mech. Lett. 48, 101438.

Schindelin, J., Arganda-Carreras, I., Frise, E., Kaynig, V., Longair, M., Pietzsch, T., Preibisch, S., Rueden, C., Saalfeld, S., Schmid, B., et al. (2012). Fiji: An open-source platform for biological-image analysis. Nat. Methods 9, 676–682.

Spurlin, J. W., Siedlik, M. J., Nerger, B. A., Pang, M.-F., Jayaraman, S., Zhang, R. and Nelson, C. M. (2019). Mesenchymal proteases and tissue fluidity remodel the extracellular matrix during airway epithelial branching in the embryonic avian lung. Development 146, dev175257.

Stancil, I. T., Michalski, J. E., Davis-Hall, D., Chu, H. W., Park, J.-A., Magin, C. M., Yang, I. V., Smith, B. J., Dobrinskikh, E. and Schwartz, D. A. (2021). Pulmonary fibrosis distal airway epithelia are dynamically and structurally dysfunctional. Nat. Commun. 12,.

Stancil, I. T., Michalski, J. E., Hennessy, C. E., Hatakka, K. L., Yang, I. V, Kurche, J. S., Rincon, M. and Schwartz, D. A. (2022). Interleukin-6-dependent epithelial fluidization initiates fibrotic lung remodeling. Sci. Transl. Med. 14, eabo5254.

Tadokoro, T., Gao, X., Hong, C. C., Hotten, D. and Hogan, B. L. M. M. (2016). BMP signaling and cellular dynamics during regeneration of airway epithelium from basal progenitors.Dev. 143, 764–773.

Tambe, D. T., Corey Hardin, C., Angelini, T. E., Rajendran, K., Park, C. Y., Serra-Picamal, X., Zhou, E. H., Zaman, M. H., Butler, J. P., Weitz, D. A., et al. (2011). Collective cell guidance by cooperative intercellular forces. Nat. Mater. 10, 469–475.

Tambe, D. T., Croutelle, U., Trepat, X., Park, C. Y., Kim, J. H., Millet, E., Butler, J. P. and Fredberg, J. J. (2013). Monolayer Stress Microscopy: Limitations, Artifacts, and Accuracy of Recovered Intercellular Stresses. PLoS One 8,.

Thomason, H. A., Scothern, A., McHarg, S. and Garrod, D. R. (2010). Desmosomes: Adhesive strength and signalling in health and disease. Biochem. J. 429, 419–433.

Tlili, S., Gauquelin, E., Li, B., Cardoso, O., Ladoux, B., Ayari, H. D. and Graner, F. (2018). Collective cell migration without proliferation: Density determines cell velocity and wave velocity. R. Soc. Open Sci. 5,.

Trepat, X., Wasserman, M. R., Angelini, T. E., Millet, E., Weitz, D. A., Butler, J. P. and Fredberg, J. J. (2009). Physical forces during collective cell migration. Nat. Phys. 5, 426–430.

Tschumperlin, D. J., Shively, J. D., Kikuchi, T. and Drazen, J. M. (2003). Mechanical stress triggers selective release of fibrotic mediators from bronchial epithelium. Am. J. Respir. Cell Mol. Biol. 28, 142–149.

Tschumperlin, D. J., Dai, G., Maly, I. V., Kikuchi, T., Laiho, L. H., McVittie, A. K., Haley, K. J., Lilly, C. M., So, P. T. C., Lauffenburger, D. A., et al. (2004). Mechanotransduction through growth-factor shedding into the extracellular space. Nature 429, 83–86.

Varner, V. D. and Nelson, C. M. (2014). Cellular and physical mechanisms of branching morphogenesis. Dev. 141, 2750–2759.

Vignaud, T., Copos, C., Leterrier, C., Toro-Nahuelpan, M., Tseng, Q., Mahamid, J., Blanchoin, L., Mogilner, A., Théry, M. and Kurzawa, L. (2020). Stress fibres are embedded in a contractile cortical network. Nat. Mater.

Vishwakarma, M., Thurakkal, B., Spatz, J. P. and Das, T. (2020). Dynamic heterogeneity influences the leader-follower dynamics during epithelial wound closure. Philos. Trans. R. Soc. Lond. B. Biol. Sci. 375, 20190391.

Walcott, S. and Sun, S. X. (2010). A mechanical model of actin stress fiber formation and substrate elasticity sensing in adherent cells. Proc. Natl. Acad. Sci. U. S. A. 107, 7757–7762.

Zhang, Z., Xia, S. and Kanchanawong, P. (2017). An integrated enhancement and reconstruction strategy for the quantitative extraction of actin stress fibers from fluorescence micrographs. BMC Bioinformatics 18, 1–14.

